# Parasitoid pressures and silence evolution

**DOI:** 10.1101/2022.08.12.503800

**Authors:** Megha R. Suswaram, Justin D. Yeakel, Chaitanya S. Gokhale

**Affiliations:** Quantitative Systems Biology, University of California Merced; Life and Environmental Sciences, University of California Merced; Research Group for Theoretical Models of Eco-evolutionary Dynamics, Department of Evolutionary Theory, Max Planck Institute for Evolutionary Biology

**Keywords:** acoustic signal, singing crickets, parasitism, reproduction, eco-evolutionary dynamics

## Abstract

Acoustic signals used by organisms to attract mates are known to attract parasitoid flies. The parasitoid flies lay their eggs inside the host signaler, eventually killing the host. We build a host-parasitoid acoustic model to investigate the effect of parasitoid flies on the signalling host’s eco-evolutionary dynamics. We used field crickets as a system to build the framework of the model. We explore how the sex ratio and the female parasitoid fecundity impact the evolution of the acoustic signal and population density of the signalling hosts. We also explore the stability of the host populations with an increase in parasitoid load. We find that up to a threshold value, an increase in parasitoid load leads to a thriving yet silent host population. Consistent with field observations, we show how this emergence of silence as an evolutionary strategy is immediate. Our results show that a drastic increase in the parasitoid load can rapidly push the signalling host population towards instability and extinction.

## Introduction

Acoustic signalling is the primary mode of communication shared by roughly 8.7 million species ranging from arthropods to mammals [1], inhabiting terrestrial and marine environments. Acoustically signalling species are incredibly diverse, from crickets, anurans, and birds to several marine organisms. The reproductive fitness can be attributed to the mating success of the individual signalers in many acoustically signalling species [2–4]. A consistent life-history trade-off is the one between reproduction and survival citekolluru1999effects. This trade-off determines the evolution of the signalling trait whether visual [5], chemical [6] or indeed acoustic acoustic [7, 8]. Signalling populations evolve to amplify or diminish their conspicuousness based on their natural chorus and other environmental pressures [9–12]. Many organisms solely use acoustic signals to secure mates. In most of these systems, the acoustic signals transmit mating opportunities with the females approaching the calling males to copulate [13]. Females cue into the signal, with some species performing phonotaxis, and locate the male. The females assess the song to assess the quality of the male [14].

Conspicuous sexual signallers, however, also garner unwanted attention. The signals risk attracting potential predators, and exploiters [15]. Male field crickets (*Gryllus campestris*) use stridulation to produce the chirp by rubbing their front wings together against the underneath of their wing, called the scraper [16]. The parasitoid flies (*Euphasiopteryx ochracea*) locate the signalling males by eavesdropping on their song by cuing in on the sound produced by the stridulation [17, 18]. The female fly then places larvae on the cricket that then burrow into the cricket’s body cavity [19]. They develop there for seven to ten days before emerging. Some parasitoid females show specificity to particular host species [15]. Studies have shown that males switch their mate securing strategies and resort to alternative mating strategies when there is an increased risk of parasitism or can even lose the signalling ability altogether [20]. Both polymorphism and plasticity has been observed in cricket populations [21, 22]. With increased parasitoid densities, male singing crickets have evolved to become silent [23]. The males develop flat wings which are incapable of producing a song. The silent males still have a chance at reproduction, as they become satellite males of the few signalling males in the population. They steal the mates from these signalling males. Additionally, added ecological pressures, compound with parasitoid population density, sex ratio, and fecundity to change the course of host signal evolution [24]. Furthermore, population dynamics create a feedback process that controls the reproduction and mortality rates, thus changing the evolutionary trajectory of sexual signals [25]. As a culmination of all the confounding factors, the evolutionary loss of the signalling traits can be remarkably rapid, in less than twenty generations, where the population becomes largely silent with a few signalers and many satellite males [23, 26].

Numerous studies focus on the evolutionary significance of parasitoid exploiters and their acoustically signalling insect hosts, such as the above-described crickets [27]. However, a quantitative estimate of the exact parasitoid densities, the proportion of parasitoid females, and other environmental pressures influencing the speed of signal loss are lacking. The existing host-parasitoid models predict the population dynamics as a version of the classical predator-prey cycles. We develop a theoretical model specific to such an extraordinary host-parasitoid system by incorporating reproductive costs and benefits of the acoustic signal.

Our mechanistic eco-evolutionary model goes beyond classical evolutionary game-theoretic reasoning that invokes negative frequency dependence. Specifically, we focus on the parasitoid sex-ratio and the parasitoid fecundity’s influence on the acoustic signal evolution and the host population density dynamics. We find a threshold parasitoid load at which there is a rapid evolutionary transition from conspicuousness to silence in the hosts. We also find that an increase in the parasitoid load decreases the stability of the host population, and the population dynamics become cyclic, chaotic and eventually go extinct. Our model will thus provide insights into the fundamental mechanisms that affect the evolution of acoustic signals in the presence of parasitoids.

## Model

We begin by recapitulating an existing host-parasitoid insect population model [28]. The population size of the host at a given time *t* is *H_t_*, and the population size of the parasitoid is *P_t_*. The proportion of parasitoid females in the population is *q*. Simultaneously, *a* is the search efficiency of the parasitoid and *F_max_* is the maximum fecundity of the parasitoid. The reproductive rate of the host is represented by *r*. This model assumes that there are i) *q* proportion of parasitoid females, ii) a parasitoid female can examine area *α* (“area of discovery”) during its lifetime, and iii) there is a maximum parasitoid fecundity, *F_max_*. The population dynamics of the host and parasitoid is given by:

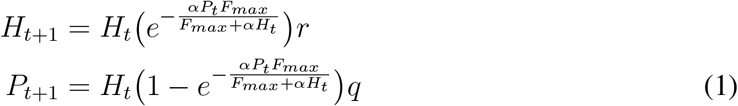

Ecologists have extensively used this particular approach to study parasitised insect populations. Host-parasitoid theoretical models typically generate oscillating populations of increasing amplitude and are by themselves unstable [29–31]. However, this does not accurately represent what happens in nature. In nature, additional ecological processes like intraspecific competition and spatial heterogeneity can partially or completely stabilise the system. The model developed by Rogers provides a realistic depiction compared to Thompson’s 1922 model [28] and Nicholson and Bailey’s 1935 model [32]. It is a further development of Holling’s disc equation [33] including realised fecundity instead of a potential fecundity [28].

### Reproductive fitness of the host based on acoustic trait

We assume that the signalling host’s acoustic trait is represented by a single acoustic character, the syllable rate. While we have considered syllable rate for building the model, it can be any feature of the acoustic signal like amplitude, frequency or intensity. We assume that the syllable rate, *z*, varies from 0 to 100 units in time. A chorus is formed when many individuals signal together with varying syllable rates. The chorus will thus have a mean syllable rate, 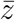. A low syllable rate is when an individual has 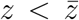. A silent individual who does not signal is represented by *z* = 0, and a high syllable rate when 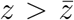 indicates a conspicuous signaler. Given an environment devoid of acoustic interference, we can assume that the mean syllable rate, 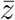, sets the reproductive fitness standard for all calling individuals. If the syllable rate of a calling individual is higher than that of the chorus mean, then the individual stands out from the overall population, is conspicuous, and can be easily distinguished by listening mates. Therefore, it has a higher chance of securing more mates. We can model this as an individual caller, whose syllable rate is higher than that of the chorus’ mean, 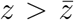, having a higher reproductive fitness component. While we assume that securing more mates increases the reproductive component of fitness, there is a maximum reproductive reward within a given time frame. The reproductive fitness with maximum reproductive reward is denoted by *r_max_*. Hence, even when the individual’s syllable reaches maximum conspicuousness, the reproductive fitness component saturates at *r_max_* (Fig. 1).

**Figure 1:**
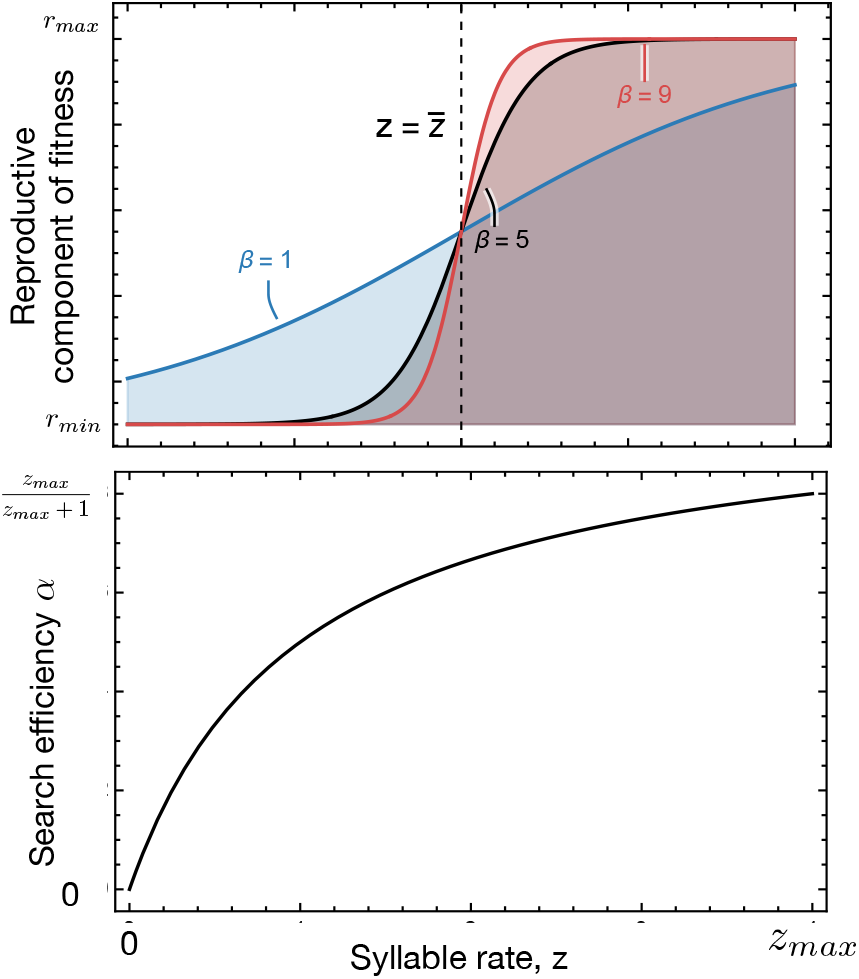
Reproductive component of fitness function of an individual host Eq. (2), Search efficiency (Eq. (3)) of a parasitoid as a function of mean chorus syllable rate. The reproductive component of fitness is dependent on how far the syllable rate of the individual, *z* is from the population chorus mean 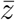, and the phonotactic selectivity, *β*. *r_min_* and *r_m_ax* are the minimum and maximum reproductive rewards respectively. With high phonotactic selectivity there is a larger benefit in reproductive component of fitness. The search efficiency of the parasitoid, *α*, increases with increase in host syllable rate and saturates at a high syllable rates.

Similarly, suppose the syllable rate of a calling individual is lower than that of the chorus’ mean. In that case, the chorus deafens the individual’s signal, and therefore it has a lower chance of securing mates. So, we can model an individual whose syllable rate is lower than the chorus’ mean, 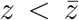, with a lower reproductive fitness component. Nevertheless, we also assume that even if a caller is silent (a non-calling individual), it can encounter a mate by random chance, with its movement within the habitat. This gives the silent individual a minimum reproductive reward value, *r_min_*. By *β* we denote the sensitivity of the reproductive curve, considered to be the phonotactic selectivity of the receiver citegerhardt2008phonotactic (Fig. 1). The reproductive component of fitness of an individual signaler is therefore given by:

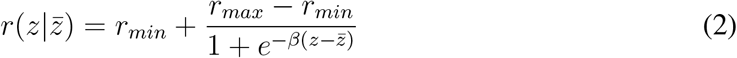

### Search efficiency of the parasitoid based on acoustic trait

We then modelled the search efficiency with a type II functional response. Hence, as the syllable rate increases, the more conspicuous the individual is, the parasitoid can better find the signaller. After a certain threshold of the syllable rate, the search efficiency of the parasitoid saturates (Fig. 1). The search efficiency *α* is,

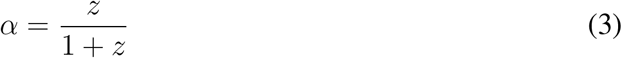

### Density dependence

We add density dependence to the insect population host-parasitoid framework. This entails replacing the host’s reproduction and search efficiency of the parasitoid with the newly formulated acoustic character-based modifications. By substituting Eq. (2) and q. (3) in Eq. (1), and adding density dependence with *K* being the carrying capacity of the habitat, we get:

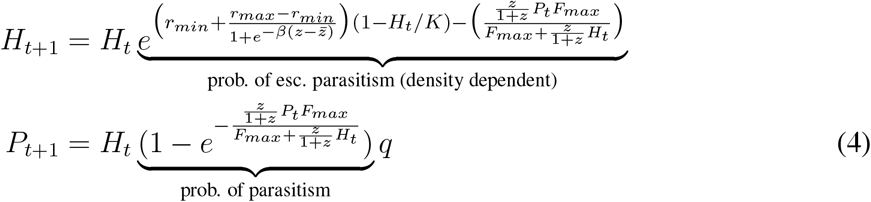

### Simulation of the eco-evolutionary dynamics

Because an analytical solution for the average fitness of the population 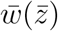 is intractable, we track changes in population abundance and the trait distributions of a cricket population with an individual-based model. As such, we numerically track the evolution of the full trait distribution of *z*, denoted by *f* (*z*), over time, in addition to population size *N* (*t*). Each time step represents a generation where all adults are assumed to die at the end of each time step such that generations are non-overlapping. Such dynamics are generally the case for cricket populations [34]; however, this assumption would not hold for many other acoustically signalling organisms, including most bird species. Offspring inherit trait values from parents with variability *σ* such that,

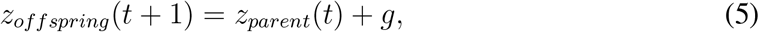

where *g* ~ N(0, *ϕ*), and we set *ϕ* = 1. The number offspring for each individual *i* with trait *z_i_* is determined by its fitness with respect to the chorus mean, 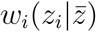, where the sum across reproducing individuals determines the future population size *N* (*t* + 1), such that

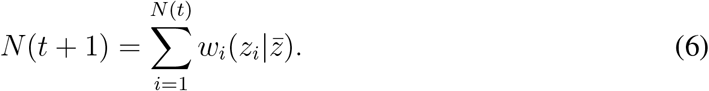

We initiate the host population to be conspicuous 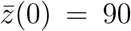 with a standard deviation of *σ* = 10, and simulate dynamics over the course of 100 generations. We confirmed 100 generations to be adequate for calculating steady-state conditions. We assumed steady state when the population density did not change more than *ϵ* = 0.00001 for more than 50 timesteps. We set *r_min_* = 0.01, *r_max_* = 2 *β* = 0.5, and *K* = 1000. The initial host population is represented in Fig. 2 (*t* = 1).

**Figure 2:**
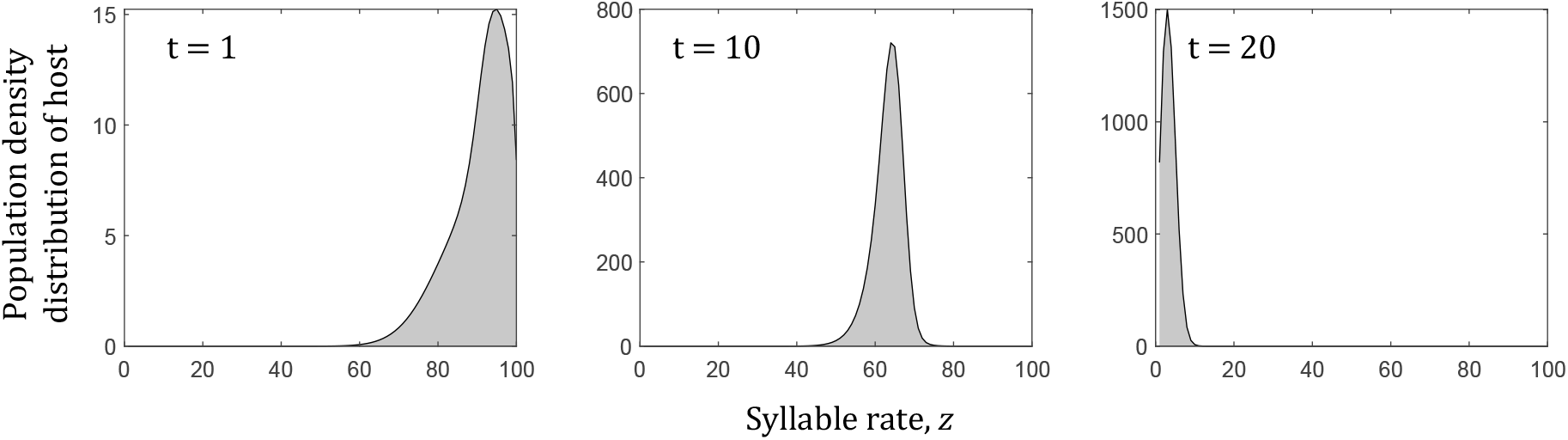
Initial distribution of the host population. With increasing generations, the host population evolves from a conspicuous chorus mean to a silent chorus mean within 20 generations. We see the evolution of silence within 20 generations in a population that has been attacked by parasitoids.The silent population has a higher population density. *H*(0) = 500, 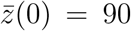, *σ* = 10, *r_min_* = 0.01, *r_max_* = 2 *β* = 0.5, and *K* = 1000.

## Results

### Quick transition from conspicuousness to silence

#### Female biased sex-ratio

As the proportion of parasitoid females increases, the conspicuously signalling population quickly evolves into silence. The reproductive fitness of the host individual is influenced by the chorus mean, 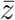. The conspicuous individuals’ fitness is initially high, resulting in reproductive gains. As the parasitoid females attack the conspicuous host individuals, mortality increases and the fitness of conspicuous individuals is reduced. In contrast, the silent host individuals escape parasitoid attack, reproduce, and increase the proportion of silent individuals in the host population in the next generation, reducing the chorus mean. With an increasing proportion of parasitoid females, there are more attacks on the conspicuous hosts, and the evolution of silence is favoured (Fig. 3). This evolutionary transition is rapid, and it usually happens within ten generations in our study. The silent individuals survive and reproduce, increasing the population density to the carrying capacity (Fig. 3). Beyond a specific female parasitoid load, all the host individuals die, and the population goes extinct (Fig. 3). Thus a female-biased sex ratio of parasitoids influences the signalling trait evolution and drives population extinction of the host.

**Figure 3:**
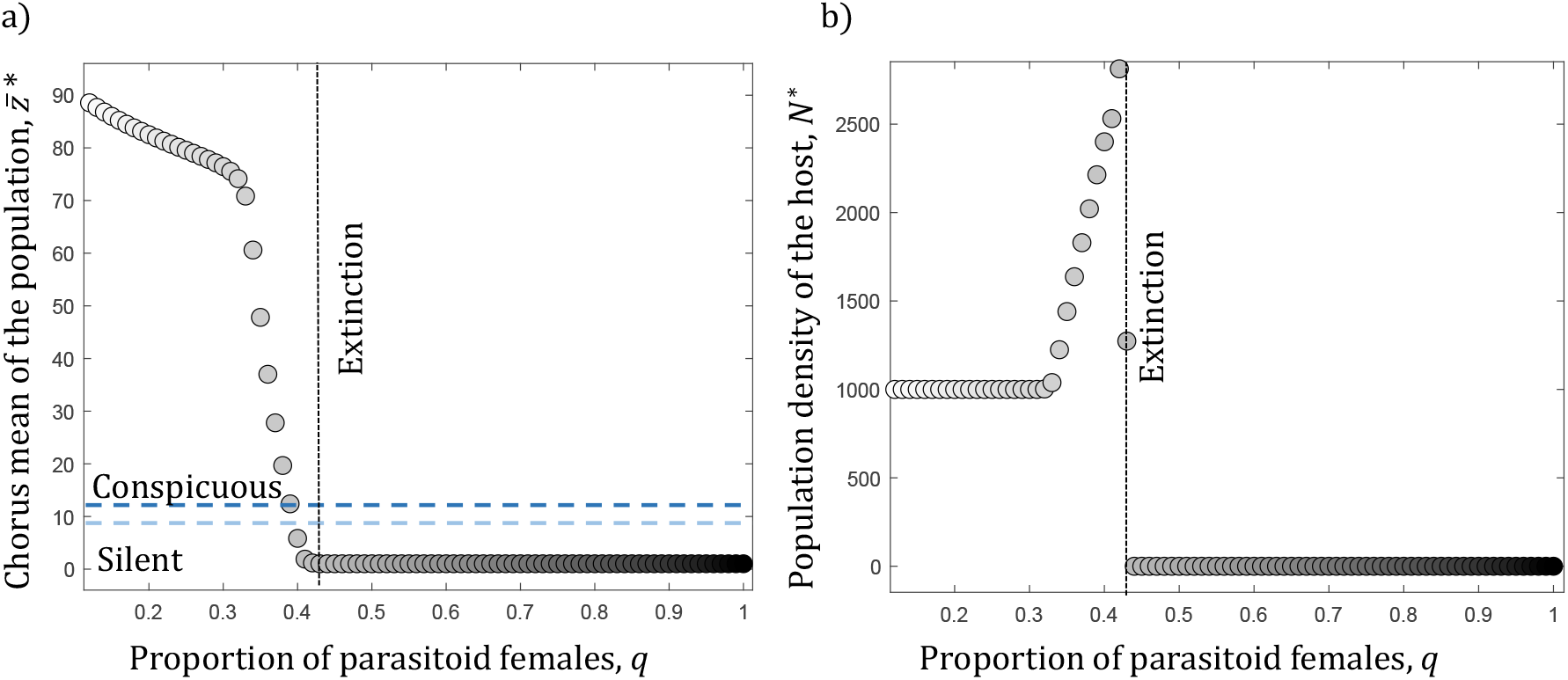
Quick transition from conspicuous to silence. a) Change in chorus mean of the host population represented by circle shapes. b) Change in population size of the host population. A more female-biased sex ratio in parasitoid populations results in a host population that is silent and large. Beyond a threshold of female-biased population in the parasitoids, the host population goes extinct. *H*(0) = 500, 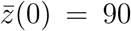, *σ* = 10, *r_min_* = 0.01, *r_max_* = 2, *β* = 0.5, *F_max_* = 3 and *K* = 1000.

#### Parasitoid fecundity influences the signalling trait evolution and population dynamics of the host

As the maximum number of viable offspring of the parasitoid females increases, the conspicuously signalling population abruptly transitions to silence (Fig. 4). The increase in parasitoid fecundity increases their population density in the next generation. This results in more number of attacks on conspicuous hosts. The conspicuous individuals die, and the proportion of silent individuals increases in the population, driving the chorus mean to silence. This evolutionary transition of the host is rapid and occurs within ten generations. As we further increase the fecundity of parasitoids, the population goes extinct (Fig. 4).

**Figure 4:**
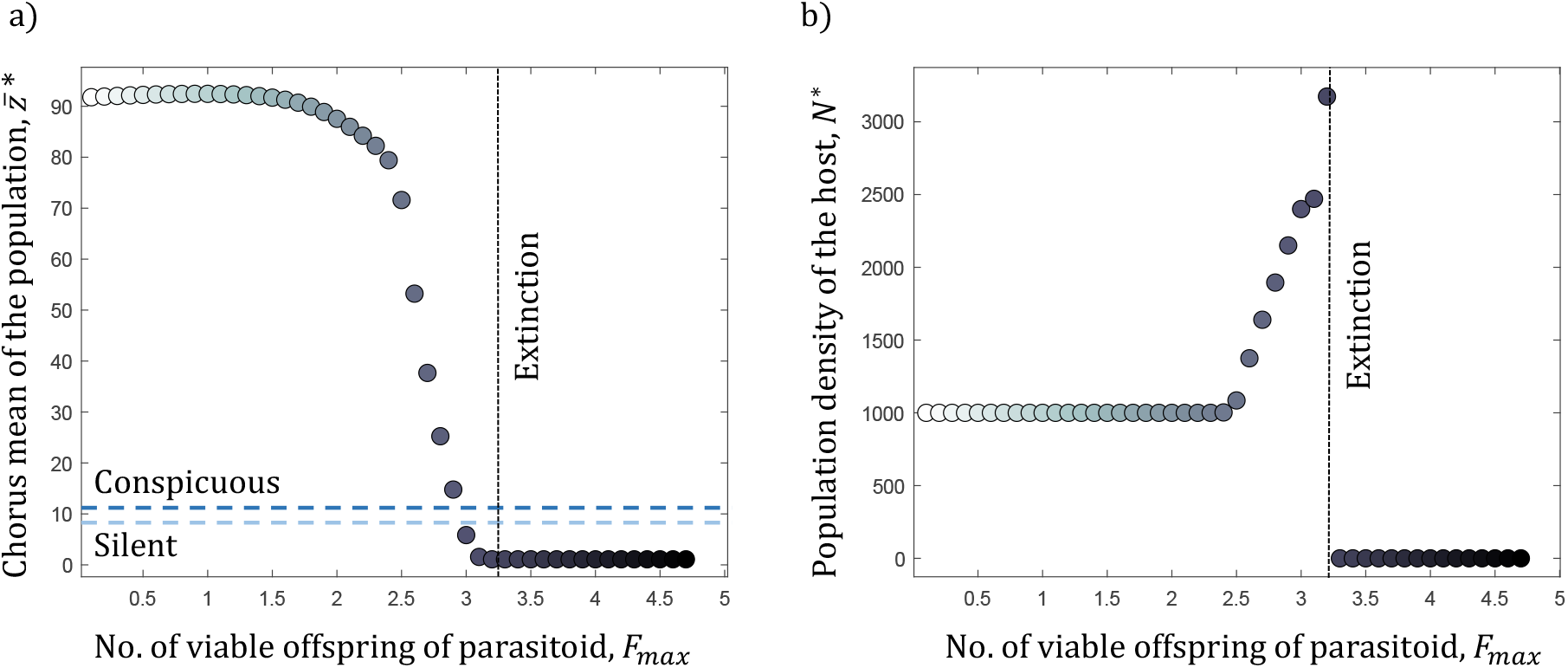
Quick transition from conspicuous to silence. a) Change in chorus mean of the host population. b) Change in population size of the host population. Increase in the fecundity of parasitoid females results in a host population that is silent and large. Beyond a threshold of fecundity of the female parasitoids, the host population goes extinct. *H*(0) = 500, 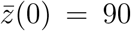, *σ* = 10, *r_min_* = 0.01, *r_max_* = 2, *β* = 0.5, *q* = 0.4 and *K* = 1000.

### Stability of the system and bifurcation analysis

The reproductive incentive to signal is given by the difference in the maximum reproductive fitness *r_max_* and the minimum reproductive reward *r_min_*. Keeping the reproductive reward constant at a low value, *r_min_* = 0.01, as we increase the maximum reproductive fitness, we increase the reproductive incentive. With the increase in reproductive incentive to signal, multiple population steady-state values emerge, such that the host population enters cyclic oscillations. We first see period-doubling in the host population density as the reproductive incentive increases. With a high reproductive incentive, the population size is similarly high. The costs of overpopulation are then determined by the carrying capacity, driving the population extinct. The stability of the system and the start of period-doubling changes with parasitoid load. As we increase the parasitoid load (increasing fecundity or the proportion of females), the populations go extinct Fig. 5.

**Figure 5:**
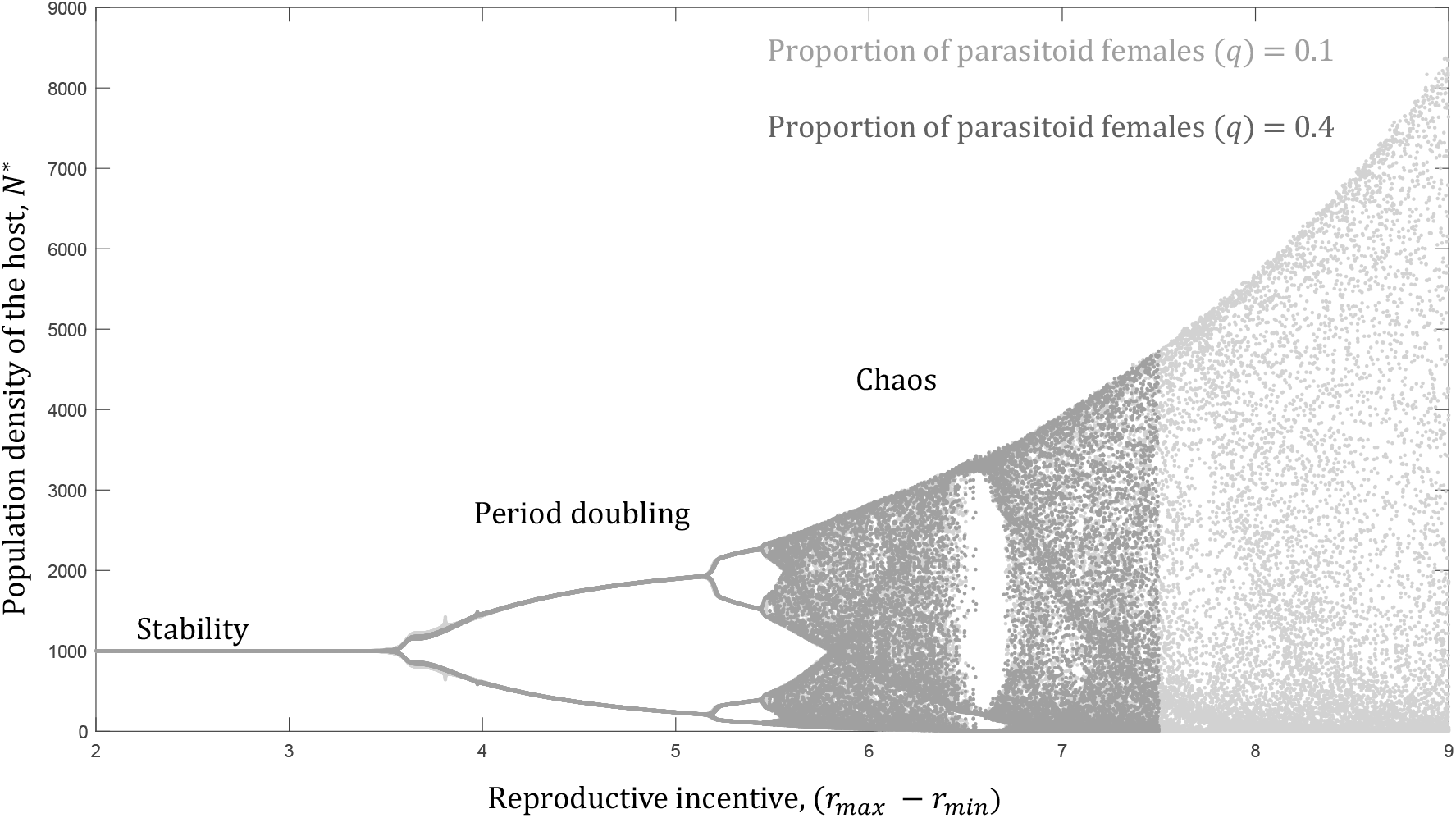
Bifurcation diagram of the host system. Outcome of host population when parasitoid population is male-biased shown in light grey and female-biased shown in dark grey. A female-biased parasitoid population causes the host population to be less stable and drive it towards extinction sooner.

We also plotted the chorus mean and the size of the host population for a varying sex ratio and fecundity of the parasitoid (Fig. 6). When the parasitoid population is more male or female biased, and the fecundity of the parasitoid is low, the host population remains conspicuous. If the sex ratio relatively is male-biased, but there is high fecundity, then the population is polymorphic with a mix of singing and silence. The host population goes extinct with a female-biased sex ratio and high fecundity (Table. 1). The system is less stable and pushed towards chaos for a female-biased parasitoid population with high fecundity and the population is subsequently driven to extinction.

**Figure 6:**
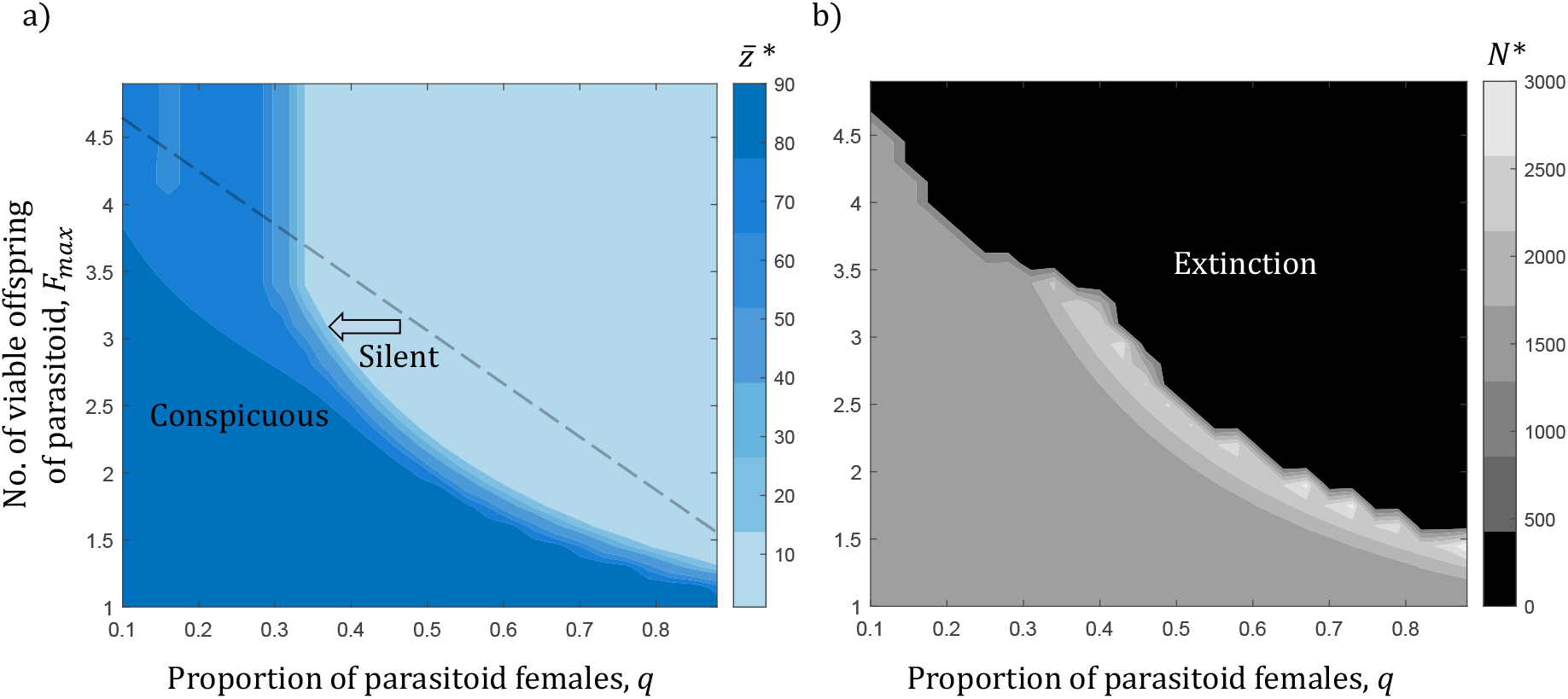
Stability analysis of the host system a) Chorus mean of the host population. Dark blue represents a conspicuous population and light blue represents a silent population. The blue transparent dashed line shows approximate extinction in population. Anything above the line is extinction and not silence. b) Population size of the host population. A high parasitoid load, pushes the host population to silence with a high population density. Beyond, a certain threshold of parasitoid load, the population enters cycles, followed by chaos, and then goes extinct. *H*(0) = 500, 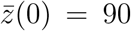, *σ* = 10, *r_min_* = 0.01, *r_max_* = 2, *β* = 0.5, and *K* = 1000.

**Table 1:** Effect of parasitoid condition on the host population

| Parasitoid Condition |  | Host Outcome |  |
| --- | --- | --- | --- |
| Sex-ratio | Fecundity | Chorus | Population density |
| Male-biased | Low | Conspicuous | High |
| Male-biased | High | Intermediate | Intermediate |
| Female-biased | Low | Conspicuous | High |
| Female-biased | High | Silent | Low |

## Discussion

Acoustic signal evolution can result from diversification of the signal [35], amplification of the existing signal to a conspicuous population citemhatre2016stay, diminishing of a signal into a silent population [36] and, loss of signal by a morphological change in the signal producing organ [23], loss of preference for conspicuous signalers from the receiver [37]. This evolution of the acoustic signal can be a result of population dynamics [38, 39], and environmental pressures [40]. Parasitism is one of the major drivers of diminishing acoustic signals in various signalling species [41]. Several empirical studies have shown that with an increase in parasitoid load, the acoustic signal is diminished and lost within a few generations [25, 42, 43] Various aspects of the effect of parasitoids on host signalling traits and population sizes have been documented [41, 44]. As discussed previously, the existing host-parasitoid models do not predict the host’s trait evolution and only focus on the population densities [28, 31, 32]. We modelled the parasitoid load, its impacts on the host population densities, and their signalling trait evolution. We have elucidated the critical components of the parasitoid load that affect the host population and identify the threshold load at which the host population responds by the loss of the signal and evolution of silence.

Our results show that the sex ratio and the fecundity of the parasitoid population play a key role in the switch from conspicuousness to silence. Specifically, the sex ratio and fecundity in the parasitoid population drive the signalling trait evolution, which in turn influences host population dynamics and stability (Fig. 3, Fig. 4, and Fig. 5). The evolution from conspicuousness to silence occurs within ten generations. Empirical studies corroborate these results. Among field crickets, it is known that signalling males are more affected by the parasitoid fly of the genus *Ormia* [18]. Female parasitoid flies find a specific cricket host depositing their parasitoid offspring. The offspring then grow and feed within the host. Within ten days, the fully grown larvae kill the host as they emerge [45]. Therefore one would estimate that the sex ratio in parasitoid populations and the offspring size play vital roles in shaping the host’s survival and signalling behaviour. Our results show that the increasing female-biased sex ratio and fecundity quickly drive the populations to evolve silence. Zuk *et al.* (2006) showed that on the Hawaiian island of Kauai, where there was high parasitoid density, most of the population evolved silence through selection for a flatwing morphology that rendered the males incapable of generating an acoustic signal within twenty generations. To reproduce, they used alternative mating strategies such as increased locomotive behaviour to encounter females randomly [44]. The resulting rapid and short transition period towards silence conforms with the results of our model.

A high reproductive incentive causes the host population destabilization, period doubling, chaos, and extinction (Fig. 5). Population densities of the host can be highly susceptible, and the risk of infection increases with parasitoid density [46, 47]. When subject to varying predator densities, Allee effects or parasitism, fast-growing populations are highly sensitive to initial population densities [48]. Such populations can exhibit chaotic dynamics that may promote extinction risk [49–53]. We show that the reproductive rate drives the population expansion, stability, and ultimately extinction (Fig. 5). Our results corroborate natural observations of rapidly growing populations which first thrive, reach a critical threshold, exhibit chaotic dynamics and then go extinct [49, 54, 55]. Also, rapid oscillations in population densities can lead to species extinction [56]. Such oscillations are dictated by the reproductive incentive, moulded by the parasitoid load. An increased parasitoid load pushes the system faster toward instability (Fig. 5). This observation implies a threshold beyond which parasitoid load cannot increase in nature as it drives the host population extinct.

Empirical studies have independently shown that parasitoids cause shifts in host traits [15]. Theoretical models and other empirical findings also point to the effects of parasitoids on host populations [28, 31, 32]. We outline the missing link: the connection between the parasitoid load, acoustic trait evolution and population density. One would expect the parasitoids first to change the host population density and the evolution of silence to be a response to the population change. Counter-intuitively, we show that the parasitoid load first alters the reproductive rate of the host population, causing an evolutionary signal adaptation in the host. The population density then responds to this evolutionary signal adaptation (Fig. 6).

Our model provides a general framework for organisms that use acoustic signals to secure mates and exploited by predators or parasitoids. This framework to predict the signal evolution change applies to crickets, cicadas, anurans, sparrows, etc. Further, the model can be expanded and developed to understand the effects of parasitism on host sex ratios [57]. The model could also be further developed to understand the evolution of multi-component auditory signals, anthropogenic effects, and specifically to climate change acoustic behaviour distributions [58–60]. Changes in the environment can cause significant changes in the reproductive incentive of signalers [40, 61, 62]. Changes in the environment can also shift the parasitoid load [63]. At different parasitoid loads, we have shown that drastic changes in evolutionary adaptations of signalling hosts and their population stability are possible for a given reproductive incentive. Together with environmental conditions that alter the reproductive incentive and parasitoid load, our model’s findings can provide insight into their reproduction, trait evolution and population densities. In future studies, we aim to leverage our findings to develop strategies for conserving the acoustic communicating populations under changing environmental conditions. Artificial manipulation of the parasitoid population size, sex ratios, and fecundity may provide a path forward.

## Acknowledgements

We thank Irina Birskis-Barros, Taran Rallings, and Ritwika VPS for their helpful comments and suggestions. This manuscript benefited from University of California, Merced’s School of Natural Sciences Dean Fellowship. Chaitanya S. Gokhale acknowledges funding from the Max Planck Society.

## Data Availability and Analysis

All code and preliminary plots are available on GitHub at https://github.com/meghasr92/parasitismsilence.

## Code availability

https://github.com/meghasr92/parasitismsilence.

## References

[1] Chen, Z. & Wiens, J. J. The origins of acoustic communication in vertebrates. Nature communications 11, 1–8 (2020).

[2] Gerhardt, H. C. & Huber, F. Acoustic communication in insects and anurans: common problems and diverse solutions (2003).

[3] Kumar, A. Acoustic communication in birds. Resonance 8, 44–55 (2003).

[4] Hedwig, B. Insect hearing and acoustic communication (2014).

[5] Nokelainen, O., Hegna, R. H., Reudler, J. H., Lindstedt, C. & Mappes, J. Trade-off between warning signal efficacy and mating success in the wood tiger moth. Proceedings of the Royal Society B: Biological Sciences 279, 257–265 (2012).

[6] Penn, D. & Potts, W. K. Chemical signals and parasite-mediated sexual selection. Trends in Ecology & Evolution 13, 391–396 (1998).

[7] Stevens, M.. Sensory ecology, behaviour, and evolution (Oxford University Press, 2013).

[8] Barbosa, F., Rebar, D. & Greenfield, M. D. Reproduction and immunity trade-offs constrain mating signals and nuptial gift size in a bushcricket. Behavioral Ecology 27, 109–117 (2016).

[9] Wollerman, L. & Wiley, R. H. Background noise from a natural chorus alters female discrimination of male calls in a neotropical frog. Animal Behaviour 63, 15–22 (2002).

[10] Vélez, A., Schwartz, J. J. & Bee, M. A. Anuran acoustic signal perception in noisy environments. In Animal communication and noise, 133–185 (Springer, 2013).

[11] Parris, K. M., Velik-Lord, M. & North, J. M. Frogs call at a higher pitch in traffic noise. Ecology and Society 14(2009).

[12] Slabbekoorn, H. & Peet, M. Birds sing at a higher pitch in urban noise. Nature 424, 267–267 (2003).

[13] Searcy, W. A. & Andersson, M. Sexual selection and the evolution of song. Annual Review of Ecology and Systematics 17, 507–533 (1986).

[14] Otter, K., Chruszcz, B. & Ratcliffe, L. Honest advertisement and song output during the dawn chorus of black-capped chickadees. Behavioral Ecology 8, 167–173 (1997).

[15] Zuk, M. & Kolluru, G. R. Exploitation of sexual signals by predators and parasitoids. The Quarterly Review of Biology 73, 415–438 (1998).

[16] Chapman, R. F. & Chapman, R. F.. The insects: structure and function (Cambridge university press, 1998).

[17] Cade, W. Acoustically orienting parasitoids: fly phonotaxis to cricket song. Science 190, 1312–1313 (1975).

[18] Robert, D., Amoroso, J. & Hoy, R. R. The evolutionary convergence of hearing in a parasitoid fly and its cricket host. Science 258, 1135–1137 (1992).

[19] Adamo, S., Robert, D. & Hoy, R. Effects of a tachinid parasitoid, ormia ochracea, on the behaviour and reproduction of its male and female field cricket hosts (gryllus spp). Journal of Insect Physiology 41, 269–277 (1995).

[20] Heinen-Kay, J. L. & Zuk, M. When does sexual signal exploitation lead to signal loss? Frontiers in Ecology and Evolution 7, 255 (2019).

[21] Rüppell, O. & Heinze, J. Alternative reproductive tactics in females: the case of size polymorphism in winged ant queens. Insectes Sociaux 46, 6–17 (1999).

[22] Ellers, J. & Liefting, M. Extending the integrated phenotype: Covariance and correlation in plasticity of behavioural traits. Current Opinion in Insect Science 9, 31–35 (2015).

[23] Zuk, M., Rotenberry, J. T. & Tinghitella, R. M. Silent night: adaptive disappearance of a sexual signal in a parasitized population of field crickets. Biology letters 2, 521–524 (2006).

[24] Rotenberry, J. T. & Zuk, M. Alternative reproductive tactics in context: how demography, ecology, and behavior affect male mating success. The American Naturalist 188, 582–588 (2016).

[25] Wilkins, M. R., Seddon, N. & Safran, R. J. Evolutionary divergence in acoustic signals: causes and consequences. Trends in ecology & evolution 28, 156–166 (2013).

[26] Tinghitella, R. Rapid evolutionary change in a sexual signal: genetic control of the mutation ‘flatwing’that renders male field crickets (teleogryllus oceanicus) mute. Heredity 100, 261–267 (2008).

[27] Lehmann, G. U. & Lakes-Harlan, R. Adaptive strategies in life-history of bushcrickets (orthoptera) and cicadas (homoptera) to parasitoids pressure on their acoustic communication systems—a case for sociality? Frontiers in Ecology and Evolution 7, 295 (2019).

[28] Rogers, D. Random search and insect population models. The Journal of Animal Ecology 369–383 (1972).

[29] Jansen, V. A. The dynamics of two diffusively coupled predator–prey populations. Theoretical Population Biology 59, 119–131 (2001).

[30] Briggs, C. J. & Hoopes, M. F. Stabilizing effects in spatial parasitoid–host and predator-prey models: a review. Theoretical population biology 65, 299–315 (2004).

[31] May, R. M. Stability and complexity in model ecosystems. In Stability and Complexity in Model Ecosystems (Princeton university press, 2019).

[32] Nicholson, A. J. & Bailey, V. A. The balance of animal populations.—part i. In Proceedings of the zoological society of London, vol. 105, 551–598 (Wiley Online Library, 1935).

[33] Holling, C. S. Some characteristics of simple types of predation and parasitism1. The canadian entomologist 91, 385–398 (1959).

[34] Masson, M. V. et al. Bioecological aspects of the common black field cricket, gryllus assimilis (orthoptera: Gryllidae) in the laboratory and in eucalyptus (myrtaceae) plantations. Journal of Orthoptera Research 29, 83–89 (2020).

[35] Symes, L., Ayres, M., Cowdery, C. & Costello, R. Signal diversification in oecanthus tree crickets is shaped by energetic, morphometric, and acoustic trade-offs. Evolution 69, 1518–1527 (2015).

[36] Gray, B., Bailey, N. W., Poon, M. & Zuk, M. Multimodal signal compensation: do field crickets shift sexual signal modality after the loss of acoustic communication?. Animal behaviour 93, 243–248 (2014).

[37] Bailey, N. W. & Zuk, M. Acoustic experience shapes female mate choice in field crickets. Proceedings of the Royal Society B: Biological Sciences 275, 2645–2650 (2008).

[38] von Helversen, D., von Helversen, O. & Heller, K.-G. When to give up responding acoustically in poecilimon bushcrickets: a clue to population density. Articulata 27, 57–66 (2012).

[39] Reichard, D. G. & Anderson, R. C. Why signal softly? the structure, function and evolutionary significance of low-amplitude signals. Animal Behaviour 105, 253–265 (2015).

[40] Bowen, A. E., Gurule-Small, G. A. & Tinghitella, R. M. Anthropogenic noise reduces male reproductive investment in an acoustically signaling insect. Behavioral Ecology and Sociobiology 74, 1–8 (2020).

[41] Kolluru, G. R. The effects of an acoustically-orienting parasitoid fly Ormia ochracea on reproduction in the field cricket, Teleogryllus oceanicus: a trade-off between natural and sexual selection (University of California, Riverside, 1999).

[42] Garamszegi, L. Z. Bird song and parasites. Behavioral Ecology and Sociobiology 59, 167–180 (2005).

[43] Rayner, J., Aldridge, S., Montealegre-Z, F. & Bailey, N. W. A silent orchestra: convergent song loss in hawaiian crickets is repeated, morphologically varied, and widespread. Ecology (2019).

[44] Balenger, S. L. & Zuk, M. Roaming romeos: male crickets evolving in silence show increased locomotor behaviours. Animal Behaviour 101, 213–219 (2015).

[45] Zukl, M., Simmons, L. W. & Cupp, L. Calling characteristics of parasitized and unparasitized populations of the field cricket teleogryllus oceanicus. Behavioral Ecology and Sociobiology 33, 339–343 (1993).

[46] Anderson, R. M. & May, R. M. Regulation and stability of host-parasite population interactions: I. regulatory processes. The journal of animal ecology 219–247 (1978).

[47] Steinhaus, E. A. Crowding as a possible stress factor in insect disease. Ecology 503–514 (1958).

[48] Hart, E. M. & Avilés, L. Reconstructing local population dynamics in noisy metapopulations—the role of random catastrophes and allee effects. PloS one 9, e110049 (2014).

[49] Schreiber, S. J. Allee effects, extinctions, and chaotic transients in simple population models. Theoretical population biology 64, 201–209 (2003).

[50] Hastings, A. Transients: the key to long-term ecological understanding? Trends in ecology & evolution 19, 39–45 (2004).

[51] Gokhale, C. S., Papkou, A., Traulsen, A. & Schulenburg, H. Lotka-Volterra dynamics kills the Red Queen: population size fluctuations and associated stochasticity dramatically change host-parasite coevolution. BMC Evolutionary Biology 13, 254 (2013).

[52] Schenk, H., Traulsen, A. & Gokhale, C. S. Chaotic provinces in the kingdom of the Red Queen. Journal of Theoretical Biology 431, 1–10 (2017).

[53] Hastings, A. et al. Transient phenomena in ecology. Science 361(2018).

[54] Sinha, S. & Parthasarathy, S. Unusual dynamics of extinction in a simple ecological model. Proceedings of the National Academy of Sciences 93, 1504–1508 (1996).

[55] Schreiber, S. J. Chaos and population disappearances in simple ecological models. Journal of Mathematical Biology 42, 239–260 (2001).

[56] Misra, J. & Mitra, A. Instabilities in single-species and host-parasite systems: period-doubling bifurcations and chaos. Computers & Mathematics with Applications 52, 525–538 (2006).

[57] Venkateswaran, V. R., Roth, O. & Gokhale, C. S. Consequences of combining sex-specific traits. Evolution 75, 1274–1287 (2021).

[58] Laiolo, P. The emerging significance of bioacoustics in animal species conservation. Biological conservation 143, 1635–1645 (2010).

[59] Roca, I. T. et al. Shifting song frequencies in response to anthropogenic noise: a meta-analysis on birds and anurans. Behavioral Ecology 27, 1269–1274 (2016).

[60] Desjonque`res, C. et al. Acoustic species distribution models (asdms): A framework to forecast shifts in calling behaviour under climate change. Methods in Ecology and Evolution (2022).

[61] Nabi, G. et al. The possible effects of anthropogenic acoustic pollution on marine mammals’ reproduction: an emerging threat to animal extinction. Environmental science and pollution research 25, 19338–19345 (2018).

[62] Lengagne, T. Traffic noise affects communication behaviour in a breeding anuran, hyla arborea. Biological conservation 141, 2023–2031 (2008).

[63] Phillips, J. N., Ruef, S. K., Garvin, C. M., Le, M.-L. T. & Francis, C. D. Background noise disrupts host–parasitoid interactions. Royal Society open science 6, 190867 (2019).

